# Gliding motility of the diatom *Craspedostauros australis* correlates with the intracellular movement of raphid-specific myosins

**DOI:** 10.1101/2024.03.11.584054

**Authors:** Metin G. Davutoglu, Veikko F. Geyer, Lukas Niese, Johannes R. Soltwedel, Marcelo L. Zoccoler, Robert Haase, Nils Kröger, Stefan Diez, Nicole Poulsen

## Abstract

Raphid diatoms are one of the few eukaryotes capable of gliding motility, which is remarkably fast and allows for quasi-instantaneous directional reversals. Besides other mechanistic models, it has been suggested that an actomyosin system provides the force for diatom gliding. However, *in vivo* data on the dynamics of actin and myosin in diatoms are lacking. In this study we demonstrate that the raphe-associated actin bundles required for diatom movement do not exhibit a directional turnover of subunits and thus their dynamics do not contribute directly to force generation. By phylogenomic analysis we identified four raphid diatom-specific myosins in *Craspedostauros australis* (CaMyoA-D) and investigated their *in vivo* localization and dynamics through GFP-tagging. Only CaMyoB-D but not CaMyoA exhibited coordinated movement during gliding, consistent with a role in force generation. The characterization of raphid diatom-specific myosins lays the foundation for unraveling the molecular mechanisms that underlie the gliding motility of diatoms.

## Introduction

Cell motility plays a crucial role in the life of unicellular eukaryotes inhabiting aquatic environments. It enables them to actively search for nutrients, evade predators, and locate suitable environments for reproduction and survival. While cilia and flagella-based swimming are well-known forms of motility, some eukaryotes, in particular protists, exhibit other remarkable modes of surface-attached movement, including crawling, rolling and gliding^1–4^. Gliding motility refers to cell movement on a surface without the involvement of external appendages such as cilia, flagella or pili^2,5,6^. While gliding motility is widespread in prokaryotes^5^, it is limited to certain eukaryotic groups such as algae (e.g., *Chlamydomonas*, diatoms) and Apicomplexans^7^ (e.g., *Plasmodium*, *Toxoplasma* and Gregarines). In these cases, the cytoskeleton plays a crucial role in generating a mechanical force, which is transmitted across the plasma membrane to adhesion complexes on the substratum, propelling the cell in the opposite direction to the force exerted intracellularly^2,6,8^. However, between the different groups of organisms, there are significant mechanistic differences in their gliding machineries.

Diatoms are a large group of unicellular algae with silica-based cell walls, found ubiquitously in sunlit aquatic habitats. Amongst the diatoms, only some pennate species that possess a specialized longitudinal slit in their cell wall, termed the raphe (and hence being called raphid diatoms), are capable of gliding motility. The raphe is an evolutionary adaption that allows diatoms to colonize and move on submerged surfaces, such as rocks, sand, animals and other algae^9^. Motile diatoms exhibit impressive velocities, reaching up to 35 µm·s^−1^, while navigating intricate, curved trajectories and being capable of quasi-instantaneously reversing their direction^10–13^. While models of the mechanism of diatom gliding date back to 1753, and numerous theories proposed over the years^14–17^, the molecular mechanism driving this unusual motility remains unknown.

The prevailing model relies on the concept of an actomyosin-based adhesion motility complex (AMC)^17–19^ and is based on the presence of two bundles of actin filaments positioned beneath the plasma membrane adjacent to the raphe opening (termed raphe-associated actin; RA-actin)^20^. According to the AMC model, there exisits a physical connection between the RA-actin through a continuum of biomolecules spanning the plasma membrane to the extracellular polymeric substances (EPS) secreted from the raphe, which enable adhesion and traction on the surface. Myosin motor proteins moving along the RA-actin are hypothesized to exert mechanical force in the AMC which is then transmitted to the substratum, enabling cell movement in the opposite direction of myosin movement^17–19^. So far, the most compelling evidence to support this model comes from experiments using free-floating microscopic beads. Beads that adhere to the raphe can display bi-directional movement on both stationary and gliding cells, with velocities similar to the velocity of the diatom gliding^17,21–23^. Recently, Gutiérrez-Medina and colleagues observed two distinct patterns of bead movement along the raphe: (i) smooth motion lasting several seconds at low velocities and (ii) intermittent, jerky motion with short (<100 ms) episodes of heightened velocity and acceleration^13^. These two patterns were found to be present during both uni-directional and bi-directional bead movement, and are attributed to a model in which cells alternate between smooth, sustained movement driven by molecular motors and abrupt, fast movement due to the elastic snapping of EPS strands. Although experiments utilizing actin and myosin inhibitors have demonstrated reversible inhibition of gliding motility^19^, direct evidence establishing actin and myosin as the principal force generators remains to be found.

Within the realm of eukaryotes, myosin motor proteins exhibit significant divergence, encompassing over 70 distinct classes, based on phylogenetic analysis of their motor domain sequences and domain architecture^24,25^. Recent investigations involving diatom genome and transcriptome sequencing projects have shed light on the phylogenetic relationships of myosins within this taxonomic group. Each diatom species possesses a repertoire of ten to eleven different myosins that fall into one of five distinct classes, and one or more myosins that remain unclassified and are referred to as ‘orphan myosins’^25^. To date, no diatom myosin has been functionally characterized. Recent advances in genetic engineering of the raphid diatom *Craspedostauros australis*^26–28^ have enabled new approaches to investigate the proposed role of myosins as force generators during diatom gliding. In this study, we aim to test this hypothesis by identifying raphid-specific myosins through phylogenomic analysis and by relating their intracellular dynamic behavior to cell movement.

## Results

### GFP-labeling allows live-cell imaging of the actin cytoskeleton in *C. australis*

Previously the actin cytoskeleton of raphid diatoms has been visualized exclusively in fixed cells stained with fluorescently labeled phalloidins^19,29–31^. To study the structure and dynamics of actin in living cells, we generated *C. australis* cell lines expressing N-terminally GFP-tagged actin. Screening of the resulting cell lines identified clones with a high ratio of GFP-labeled actin that were used for the precise localization of the actin cytoskeleton, and clones with lower amounts of GFP-labeled actin (‘speckled’) that were used for quantitative analysis of actin dynamics. The latter was crucial to (i) ensure that fluorescent labeling of the actin bundles is discontinuous, thereby enabling the detection of actin dynamics, and (ii) to minimize potential artifacts known to be caused by GFP-tagging of actin that could compromise its function^32,33^. Motile cells were observed in all clones irrespective of the abundance of actin-GFP, indicating that the GFP tag did not inhibit cell motility.

Confocal fluorescence microscopy confirmed the presence of two RA-actin bundles that are continuous along the longitudinal axes of the two opposite valves, which together with the girdle band, form the diatom frustule. There is a distinct separation of the two RA-actin bundles in the center of the cell, beneath the central nodule. At the cell apicies, the RA-actin bundles follow the shape of the valves (**Fig. 1a, c**) and do not extend into the girdle band region (**Fig. 1b, d**). Imaging of the cell apices revealed that the two RA-actin bundles appear to be continuous, forming narrow loops each with a diameter of ∼250 nm (**Fig. 1e, f**). In addition, a pronounced ring of actin filaments is associated with the girdle bands and an extensive network of actin was observed in the perinuclear region of the cell, which corresponds to the position of the Golgi complex (**Fig. 1d, Supplementary Fig. 1**).

**Fig 1:**
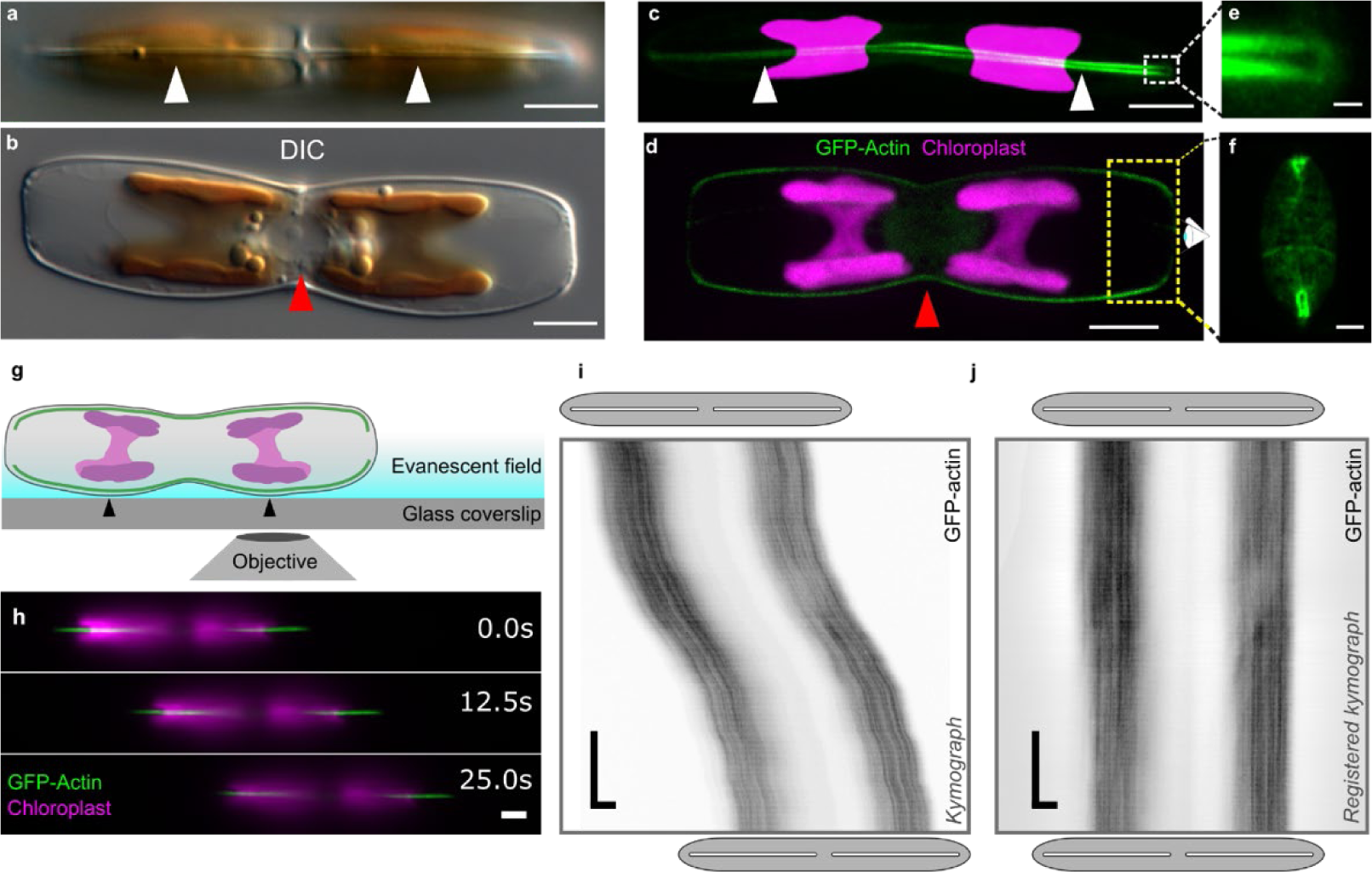
Analysis of actin structure and dynamics. **a, b** Differential interference contrast (DIC) light microscopy images of *C. australis* wild type cells in valve view (**a**, white arrowheads point to the raphe slit) and girdle view (**b**, red arrowhead points to the perinuclear region). The chloroplast has a golden-brown pigmentation. **c, d** Confocal fluorescence microscopy images of live cells embedded in agarose expressing GFP-actin. Z-projection of optical sections of the proximal raphe slit adjacent to the glass coverslip **c** (white arrowheads point to the same position as in a) and the midline section of the cell **d** (red arrowhead pointing to perinuclear actin). **e, f** Magnified loops of raphe-associated GFP-actin at the distal ends of cells embedded in agarose in raphe view (**e**, magnification of white dotted box in **c**) and in oblique view (**f,** approximate location of yellow dotted box shown in **d**, rotated 90 degrees away from the viewer, shown for a different upright cell). **g** Schematic of live cell TIRFM experimental setup. Cells adhere and glide with one set of actin bundles (green) and two chloroplasts (magenta) in the evanescent field close to the coverslip surface. Black arrowheads indicate points where the cell wall is in close proximity to the coverslip. **h** Montage generated from dual-color TIRFM images showing the position of a gliding cell at three time points. **i, j** Kymograph analysis of GFP-actin movement (in black) using raw **i** and registered **j** GFP-actin data from **h**. Gray ellipses and white slits above and below kymographs approximate the positions of the cell body and raphe openings at the beginning and end of imaging, respectively. (green – actin; magenta – chloroplast autofluorescence; scale bars: a-d, h: 5 µm, e: 0.5 µm, f: 2 µm, i, j: horizontal 5 µm, vertical 5 s)

### Actin dynamics do not contribute to the generation of force for cell gliding

To investigate the possible involvement of actin dynamics (for example via filament treadmilling) in gliding motility, we performed time-lapse dual-color total-internal-reflection fluorescence microscopy (TIRFM) on both mobile and stationary cells expressing GFP-tagged RA-actin (**Fig. 1g-j**). This technique enabled the simultaneous imaging of the GFP-tagged RA-actin and chloroplast autofluorescence in the evanescent field of TIRFM close to the coverslip (**Fig. 1g**). We developed an image analysis pipeline that utilizes tracking of chloroplast autofluorescence as a proxy for cell movement. By effectively subtracting the cell movement from the kymograph (space-time plots of the fluorescence intensities along the direction of the raphe), we were able to relate the intracellular movement of fluorescently labeled actin (or myosin, see below) with respect to cell movement (**Supplementary Fig. 2, Methods**).

A technical limitation encountered during TIRFM imaging of the GFP-labeled RA-actin bundles arises from their spatial distance from the coverslip. This distance is subject to variations as (i) the RA-actin bundles follow the curved shape of the cell wall (**Fig. 1b, d**)^27^ and (ii) the EPS layer varies in thickness during gliding motility. Consequently, it is challenging to visualize the entire length of the bundles during cell movement. Nonetheless, as the cells glided across a coverslip, several micrometer-long sections of the GFP-actin bundles were visible along the longitudinal axis of the cell (**Fig. 1h, Supplementary Fig. 3a, Supplementary Movies 1, 2**). Kymograph analysis revealed that there was no discernable net movement of the GFP labeled RA-actin relative to the cell in gliding or stationary cells within time intervals of 25-30 s (example kymographs in **Fig. 1i, j, Supplementary Fig. 3b**). These observations indicate that there is no directional turnover of actin subunits within the RA-actin bundles, and thus we proceeded with the assumption that actin dynamics do not contribute to force generation for gliding.

### Phylogenomic analysis reveals four *C. australis* myosins in a raphid-specific clade

Myosins are currently classified into 79 classes based on phylogenetic analysis of their motor domain sequence^25^. As gliding motility is restricted to raphid diatoms, we performed a phylogenomic analysis on all publicly available diatom myosin sequences to determine which myosins are specific to raphid diatoms. BLAST analysis of the *C. australis* genome assembly^27,28^ revealed twelve putative myosin sequences, a number which is similar to other sequenced diatoms (*Phaeodactylum tricornutum*: 10; *Thalassiosira pseudonana*: 11)^34^. The phylogenomic analysis was performed using the motor domains of 320 diatom myosins from 52 diatom species (**Supplementary Fig. 4**) and corroborated previous findings distinguishing five diatom myosin classes (Class 29, 47, 51, 52 and 53) and between one and four orphan, unclassified myosin sequences^24,25,34,35^.

Four of the *C. australis* myosins, which we named CaMyoA-D (all belonging to myosin Class 51), were exclusively found in raphid diatoms (**Supplementary Fig. 4**, highlighted in blue). CaMyoA-D possess canonical myosin domains (motor, neck, tail) and contain essential motor domain motifs including the purine binding loop, Walker A motif, switch 1/2 regions, actin-binding site, Src homology 1 helix, and possess variable numbers of IQ motifs in the neck region (**Fig. 2a**). The tail domains of these four myosins are relatively short (276-417 amino acids), do not contain any additional protein domains, and are predicted to contain coiled-coil regions suggesting they are dimeric *in vivo*. Based on the phylogenomic distance of these four myosins to previously described *P. tricornutum* myosins suggests homology of: CaMyoA – PtMyoG, CaMyoB/C – PtMyoC/A, CaMyoD-PtMyoE^34^.

**Fig. 2:**
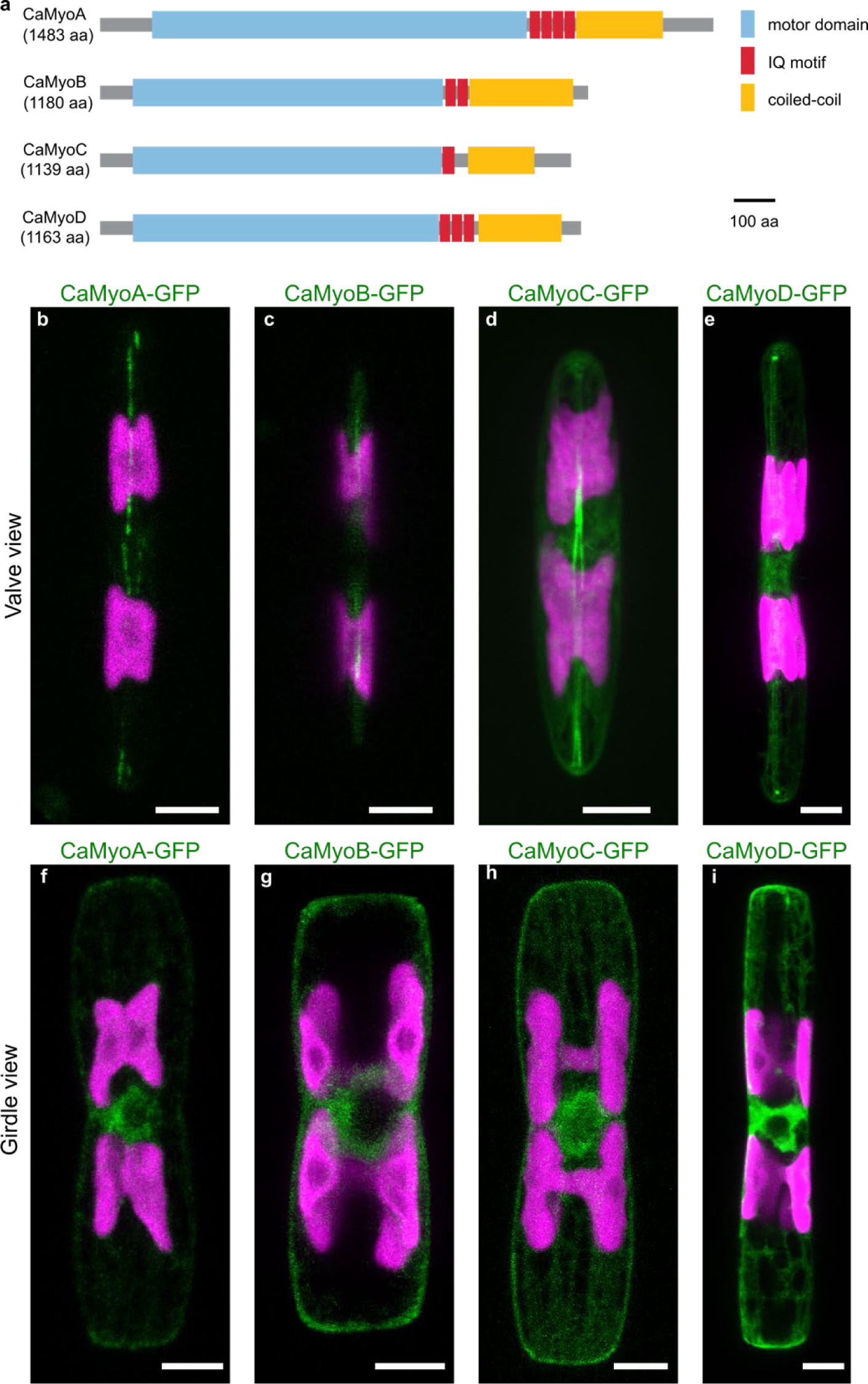
Schematic primary structures and intracellular localization of CaMyoA-D. **a** Schematic domain structure of the four myosins investigated in this study. Scale bar: 100 amino acids. **b-i** Localization of myosin-GFP fusion proteins. Each image is a maximum z-projection of the confocal fluorescence microscopy stack of the indicated GFP-tagged myosin. GFP-tagged myosin fluorescence shown in green and chloroplast autofluorescence in magenta. Scale bars: 5µm.

### CaMyoA-D are distributed along the longitudinal axis of the valve and in the perinuclear region

To explore the involvement of CaMyoA-D in generating force for gliding motility, we expressed them as C-terminally tagged GFP-fusion proteins. First, we visualized their intracellular localization by confocal fluorescence microscopy in cells embedded in agarose, where gliding motility was suppressed (**Fig. 2b-i**). All four GFP-tagged myosins (i) were localized along the longitudinal axis of each valve with varying degrees of continuity in the raphe region, (ii) exhibited a high abundance around the perinuclear region, and (iii) were distributed throughout the cytoplasm in a mesh-like network similar to the distribution of GFP-tagged actin.

CaMyoA-GFP was typically seen localized along the longitudinal axis of both valves in two interrupted linear patterns, which extended to the cell apices and like the RA-actin bundles diverged near the center of the valve where the central nodule is located. In contrast, CaMyoB-GFP was typically only observed in discrete fluorescent regions along the longitudinal axis of the valve, and it was not possible to distinguish two separate linear patterns. Based on the position of the chloroplasts, the discrete regions of CaMyoB-GFP fluorescence were close to the curved regions of the valve, and likely closer to the cover glass (see **Fig. 1g**). CaMyoC-GFP was typically seen in two dense linear patterns along the longitudinal axis of both valves. Like CaMyoA-GFP, the distance between these linear patterns was seen to diverge near the central nodule. CaMyoD-GFP was also seen in two linear patterns along the longitudinal axis both valves, but additionally showed distinct accumulations at the cell apices.

### TIRFM reveals dynamic inhomogeneities of CaMyoA-D along the longitudinal axis of the cell

To investigate the dynamic behavior of the myosins, we performed live cell dual-color time-lapse TIRFM imaging on both mobile and stationary cells with GFP fusion proteins of CaMyoA-D (**Fig. 3**). For all four myosins we observed a rather uniform distribution of GFP-fluorescence along the entire longitudinal axis of each valve, closely matching the position of the RA-actin bundles. In addition, we observed some inhomogeneities in the GFP-fluorescence signal, manifesting as individual or multiple distinct spots. Specifically, these spots were observed for (i) CaMyoA-GFP in predominantly stationary (or slow-moving) cells, (ii) CaMyoB-GFP and CaMyoC-GFP exclusively in gliding cells, and (iii) CaMyoD-GFP both in stationary and gliding cells.

**Fig. 3:**
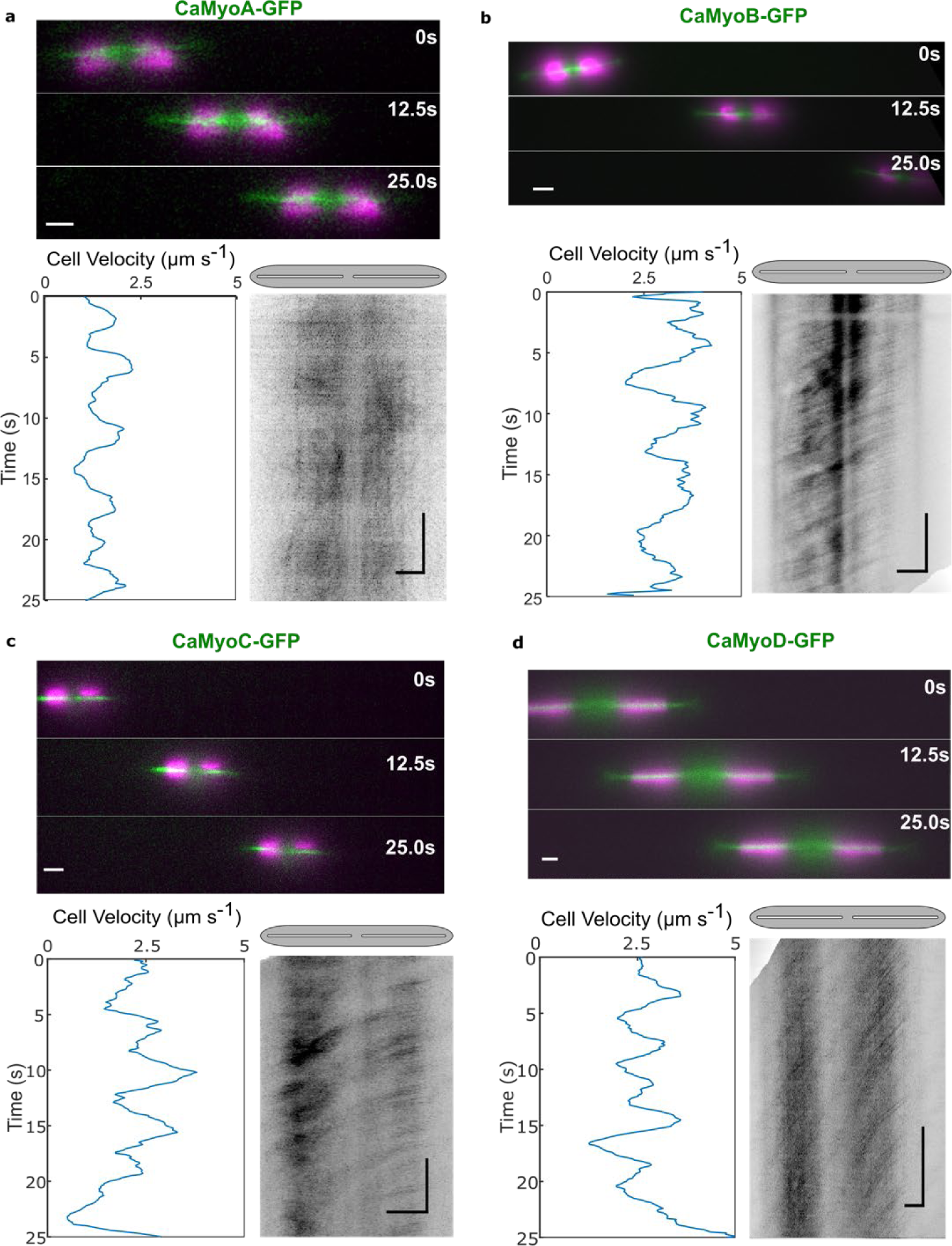
Dynamic localization of CaMyoA-D in cells during, smooth sustained gliding. Analysis of 25 s time-lapse segments of gliding cells expressing **a** CaMyoA-GFP, **b** CaMyoB-GFP, **c** CaMyoC-GFP, and **d** CaMyoD-GFP. (Upper panels) Montages showing the position of cells at 12.5 s intervals (GFP in green, chloroplast autofluorescence in magenta, scale bars: 5 µm). (Lower panels, left) Cell velocity as function of time, generated from chloroplast tracking data. (Lower panels, right) Registered kymographs generated from GFP-channel data (black) showing movement of myosins relative to the cell. Gray ellipses and white slits above kymographs approximate the positions of the cell body and raphe openings, respectively. (Scale bars: horizontal = 5 µm, vertical = 5 s)

According to the AMC hypothesis, we would expect a myosin contributing to cell motility (i) to be active on the RA-actin bundles, (ii) exhibit little to no movement relative to the substratum, and (iii) move towards the trailing end of the moving cell. To investigate which myosins show such behavior, we analyzed the dynamic behaviour of individual spots of myosin-GFP fusion proteins in continuously moving cells. CaMyoA-GFP spots occasionally showed slow, non-coordinated movement (**Fig. 3a, Supplementary Movie 3**). Spots of CaMyoB-GFP, CaMyoC-GFP and CaMyoD-GFP were frequently observed to move persistently and in a coordinated manner towards the trailing end of the cell (**Fig. 3b-d, Supplementary Movies 4-6**). The trajectories of spots of CaMyoB-GFP, CaMyoC-GFP and CaMyoD-GFP were parallel to the long axis of the valves, crossed the midpoint of the valve and often exhibited nearly constant velocities over extended time periods.

### Intracellular motility of CaMyoB-D, but not CaMyoA, correlates with cell motility

We further quantified the correlation between myosin and cell motility using our image analysis pipeline described above. We found that the majority of CaMyoA-GFP spots moved in long, slow runs in both directions within stationary and slowly-moving cells (**Supplementary Fig. 5, Supplementary Movies 7-9**). CaMyoA-GFP spots often passed one another, moving in the opposite direction and their mean velocity was typically slower than the gliding velocity of the cells (**Fig. 4**). Moreover, the spots did not strictly follow the centerline of the valve, suggesting that CaMyoA-GFP is not exclusively moving on the RA-actin bundles. In contrast, the CaMyoB-GFP, CaMyoC-GFP and CaMyoD-GFP spots moved exclusively in opposite direction to cell movement. The absolute values of the associated myosin velocities (ranging up to 12 µm s^−1^) always exceeded the cell velocities (ranging up to 4 µm s^−1^) (**Fig. 4**).

**Fig. 4:**
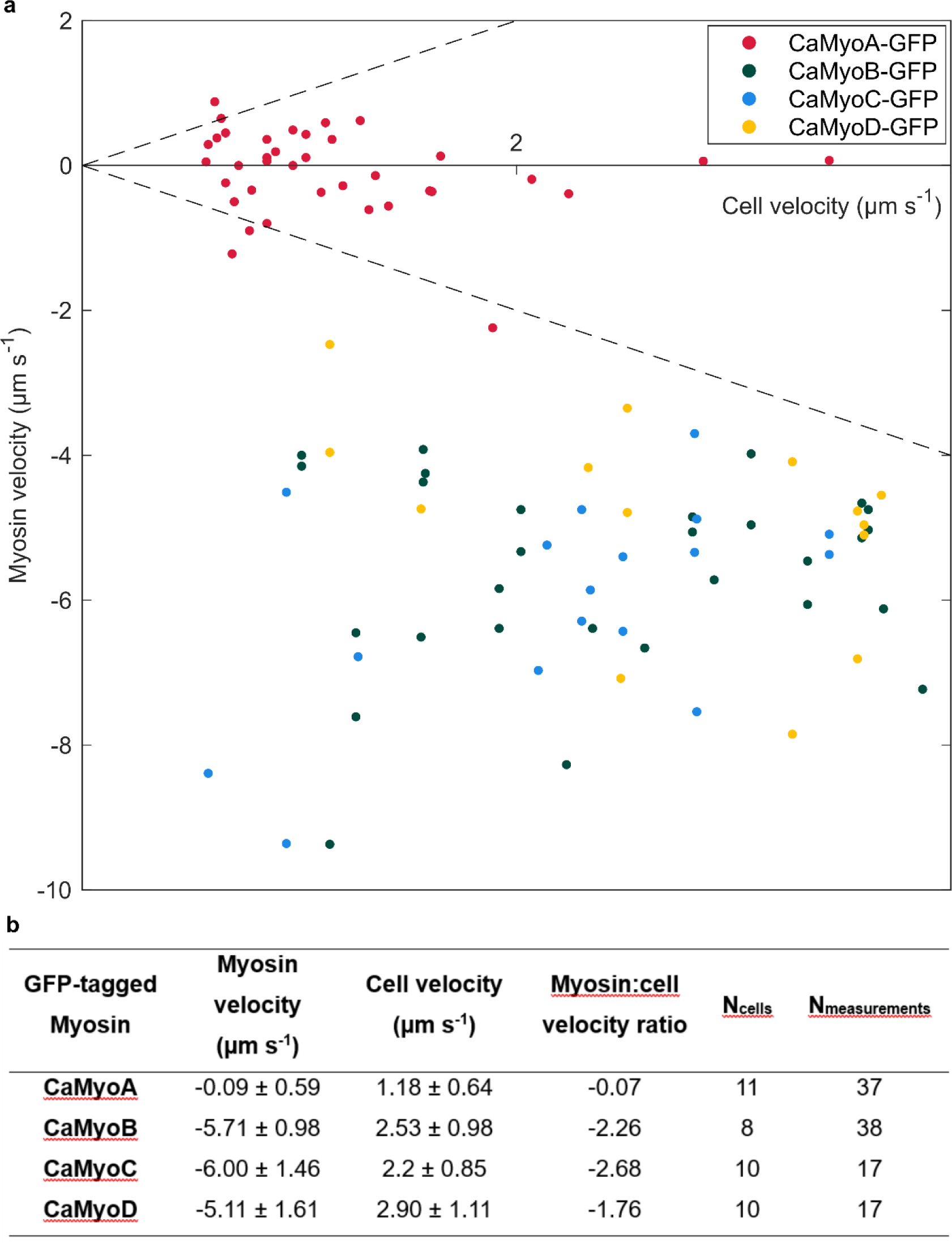
Quantitative relationship between cell velocities during smooth, sustained gliding and intracellular GFP-tagged myosin velocities. **a** Mean myosin velocity relative to the moving cells (each data point signifies one measurement in one half of the cell during a given time window, see Materials and Methods) as function of mean cell velocity (during the same time window). Dashed line indicates the equality of myosin and cell velocity. **b** Myosin velocities (mean +/- SD) and corresponding mean cell velocities (mean +/- SD) along with their ratio, as well as, number of cells (N_cells_) observed and total measurements (N_measurements_) performed.

If intracellular myosin motility were to be directly involved in propelling cell motility, one would further expect a correlation between myosin and cell velocities upon changes in the motility state of moving cells. We indeed observed such a correlation for (i) starting cells (i.e. cells transiting from a resting state to a moving state, **Fig. 5a**), (ii) stopping cells (i.e. cells transiting from a moving state to a resting state, **Fig. 5b**), and (iii) cells reversing their direction (**Fig. 5c**). In all cases, the motility of CaMyoB-GFP (which exhibited the most consistent correlative behavior among the myosins) showed fast coordinated movement when gliding started, abrupt halting when gliding stopped and quasi-instantaneous directional reversal when gliding switched direction (**Fig. 5, Supplementary Movies 10-12**). Similar behaviors were observed for CaMyoC-GFP and CaMyoD-GFP spots, though they occasionally moved in opposite directions (either towards the center or towards the apices of the cell valve) in the leading and trailing halves of the same cell (**Supplementary Fig. 6, Supplementary Movies 13-15**). The latter cases were rarely observed during smooth, sustained gliding but predominantly in slow-moving cells exhibiting frequent fluctuations in their velocity and/or their direction of cell movement.

**Fig. 5:**
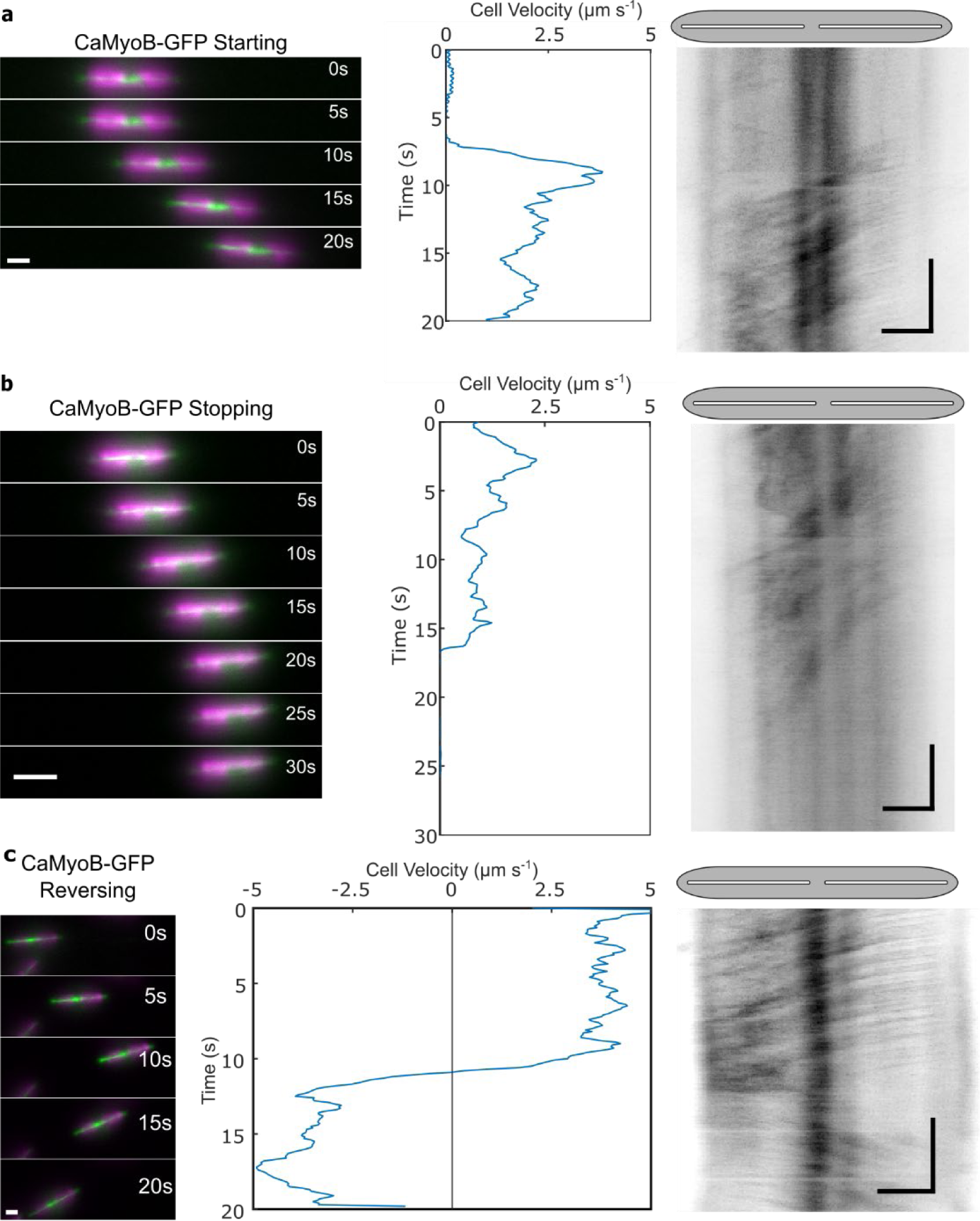
Dynamic localization of CaMyoB in cells during changes in their motility state. Analysis of 20-30 s time-lapse segments of CaMyoB-GFP expressing cells during **a** starting, **b** stopping and **c** reversing. (Left panels) Montages showing the position of cells at 5 s intervals (GFP in green, chloroplast autofluorescence in magenta, scale bars: 5 µm). (Middle panels) Cell velocity as function of time, generated from chloroplast tracking data. (Right panels) Registered kymographs generated from GFP-channel data (black) showing movement of myosins relative to the cell. Gray ellipses and white slits above kymographs approximate the positions of the cell body and raphe openings, respectively. (Scale bars: horizontal = 5 µm, vertical = 5 s)

## Discussion

In this study, we investigated the involvement of actin dynamics and four raphid-specific myosins (CaMyoA-D) in the gliding motility of the diatom *C. australis*. Our results demonstrate that diatom gliding does not rely on a direct force contribution from actin treadmilling, and is thus markedly different from the mechanisms of lamellipodia-driven crawling motility or apicomplexan gliding motility that both require actin polymerization^36,37^. Because of the rigid diatom cell wall, movement driven by actin-based cell membrane protrusions would need to extrude through the raphe as proposed in the actin pin hypothesis^16,38^. However, our actin localization experiments showed no evidence of actin-based cell membrane protrusions through the raphe and beyond the cell wall. Additionally, many diatom species have been shown to lack profilins and the Arp2/3 complex^39^, both of which are essential regulators of actin branching and nucleation required for membrane protrusions in other eukaryotes. The precise architecture of the RA-actin bundles, such as the length of individual actin filaments and their polarity within a single bundle remains unknown. Consequently, how these factors may contribute to the bi-directional diatom gliding mechanism remains an open question. Based on our observation that both cells and intracellular myosins can quasi-instantaneously reverse their direction, several possibilities emerge: (i) the two actin bundles are antiparallel and unidirectional, (ii) both actin bundles are bidirectional, or (iii) at least one of the myosins is capable of moving towards the minus-end of the actin filaments. Notably, our investigation into the RA-actin bundles revealed loop-like structures at the cell apices connecting the two actin bundles, suggesting the intriguing possibility that they may form a single continuous bundle. In this case, such a continuous track would eliminate the need for secondary transport of myosins once the end of a bundle is reached, even if the individual actin bundles are not bidirectional.

Our results demonstrate that CaMyoB-D, but not CaMyoA, engage in fast, coordinated movement in the opposite direction to cell movement. This inverse relationship between the direction of myosin movement and cell gliding aligns with the hypothesized force coupling mechanism within the AMC model^17^. Although CaMyoB-D displayed comparable kinetic behaviors, which may indicate some level of redundancy, we cannot rule out additional isoform-specific functions, which may play a role during the generation of complex trajectories, directional changes and intracellular transport.

In their natural habitat, diatoms must navigate their surroundings, respond to chemical gradients, and migrate towards or away from specific cues (e.g. light, nutrients, toxins)^40–43^. Remarkably, they accomplish these tasks with a high degree of temporal and spatial control^44^. Individual diatom species can vary their cell velocities by one order of magnitude, are capable of complex movements in circular, spiral, straight and even sometimes sigmoidal trajectories and are able to reverse their cell direction quasi-instantaneously^11,12,45^. While our study has yet to investigate the behavior of GFP-tagged CaMyoB, -C, and –D, simultaneously within a single cell line or over longer cell trajectories, it is conceivable that the distinct behaviors observed for each myosin contribute to disrupting symmetry and distributing forces along the entire raphe. This, in turn, may pave the way for changes in direction and velocity of cell movement. Consequently, by modulating the activity of a combination of myosin isoforms, diatoms may be capable of generating forces in both directions along the raphe, ultimately underpinning the intricate trajectories witnessed in the gliding diatoms.

During smooth, sustained gliding, multiple spots of each CaMyoB, CaMyoC and CaMyoD move at uniform velocities, suggesting a high degree of myosin cooperativity along the entire length of the cell. Theoretically, the force generated by a single myosin would be sufficient to overcome the viscous drag of a diatom moving in aqueous solution; according to Stoke’s law the viscous drag of a cylinder with half-sphere ends and a diameter of 6 µm moving through water at 4 µm s^−1^ is less than 1 pN. Nevertheless, recent research has shown that diatoms are capable of generating forces far greater (∼ 800 pN) than those required to propel cell movement, which is consistent with a large number of myosin motors being simultaneously involved in force generation^45^. This becomes particularly relavant when considering the natural habitat of raphid diatoms, where they must force their way through fine-grained sediments laden with obstacles, which is in stark contrast to the controlled environment of a smooth laboratory glass slide. Furthermore, when embedded within a dense biofilm matrix, diatoms must exert sufficient mechanical forces to overcome the elastic resistance of the biofilm EPS. It is therefore likely that cooperative myosin activity enables (i) the generation of force large enough to overcome obstacles and (ii) the coordination of directional forces exerted at multiple raphe openings to establish, maintain or reverse the direction of gliding. We hypothesize that the cooperative activity of myosin may be orchestrated through chemical signaling pathways (such as phosphorylation or calcium ions)^46^ or physical interactions (such as oligomerization of their tail domains)^47,48^.

We consistently observed that the velocity of cell movement was slower (up to 50%) compared to the velocity of the GFP-tagged myosins. This finding strongly suggests mechanical compliance in the motility machinery. According to the AMC model, diatom gliding requires the transmission of a force generated by the intracellular cytoskeleton across the plasma membrane and through the raphe slit to the extracellular substratum via adhesive EPS strands. Consequently, the loss in velocity, which indicates a loss in force transduction from the actomyosin complex to the substratum, could be caused by (i) inefficient force transfer through the plasma membrane and raphe slit (similar to slippery motor anchorage^49^), or (ii) the mechanical properties of the EPS strands, which exhibit high tensile strength and elasticity attributed to the presence of large modular proteins^50^. Some of these EPS strands, in particular those not connected to the AMC, may exert elastic or frictional counterforces potentially causing a dissipation of energy during force transmission^50^. Recent findings have started to unveil the molecular composition of the diatom EPS and identified two exceptionally large modular proteins in the *C. australis* adhesive trails (CaTrailin4; ∼900 kDa and Ca5609; >1 mDa)^28^, which may contribute to these counterforces, and a mucin-like protein (CaFAP1)^27^ that may contribute to reducing the frictional resistance between the cell and the substratum. However, how these components, and others, contribute to mediating interactions between the intracellular cytoskeleton, the cell surface and the substratum beneath remain to be investigated.

Occasionally, spots of CaMyoC and CaMyoD moved in a fast and coordinated manner toward the center of a gliding cell in both the leading and trailing halves. This phenomenon typically coincided with an abrupt change in the direction of cell movement or comparatively low gliding velocities (<1.5 µm·s^−1^), suggesting that these myosins can generate opposing forces in each half of the cell. Opposing forces play an important role in many cytoskeletal processes including bidirectional transport of vesicles, cell size determination, and actin ring formation in cytokinesis, which rely on similar tug-of-war systems between motors^51–53^. *In vitro* experiments have also shown that groups of molecular motors pulling in opposite directions can cause spontaneous symmetry breaking^54^ and distinct modes of slow and fast movements, as well as sharp transitions between these modes^55^. The observation of counteracting force production by both CaMyoC and CaMyoD corroborates findings from previous research, which has indirectly demonstrated the existence of such opposing forces in diatom motility^18,56^.

CaMyoA displayed a single mode of very slow, non-cooperative myosin movement without any observed coordination with overall cell movement. This movement pattern bears resemblance to the characteristic behavior of myosins engaged in intracellular transport, such as class V myosins. The high duty ratio of class V myosins enables them to move processively during force generation resulting in long, slow runs during which the myosin does not detach from actin^57,58^. Such intracellular transport may contribute to gliding by delivering vesicles containing EPS to the raphe.

Taken together, our findings suggest that the concerted action of raphid-specific myosins directly drives diatom gliding motility. Beyond these results, our work opens avenues for further exploration of myosin functions in other eukaryotes as well as for the design of smart synthetic mechanosystems inspired by biological organisms. As such, it would be intriguing to design molecular transport, manipulation and sorting devices that combine the distinct diatom properties of large forces, high velocities and rapid directionality switching. In future studies, it will be of interest to track CaMyoB, CaMyoC and CaMyoD simultaneously in the same cell. The results of such experiments will reveal whether these myosins work in concert to drive smooth, sustained gliding, or if they are capable of fulfilling this role interchangeably. Moreover, correlating the movement of intracellular myosins to the movement of extracellular particles or the deformation of the substratum will allow further quantification of the force transmission from the molecular actomyosin system to the exterior of the cell.

## Methods

### Diatom strains and culture conditions

*Craspedostauros australis* (Cox, CCMP3328) was grown in artificial seawater medium, EASW^59^ or ASW^60^ at 18 °C under constant light and intensity between 40 and 60 µmol photons·m^−2^ ·s^−1^, using a cool white light source and unless otherwise stated, maintained with a 14/10-hour light/dark cycle. All cell lines were subcultured regularly (every 7-10 days) to maintain cells in a logarithmic stage of growth. All mutant cell lines were maintained in a medium supplemented with 450 µg/mL nourseothricin (clonNAT) antibiotic (Jena Biosciences, Jena, Germany).

### Identification of myosin and actin genes in *C. australis*

To identify putative *C. australis* myosin genes a tBLASTn protein homology search using available myosin and actin proteins from *Phaeodactylum tricornutum*^34^ was performed against the *C. australis* genome assembly^27,28^. All protein sequences were validated for the presence of conserved myosin and actin domains using an Interpro search (https://www.ebi.ac.uk/interpro/).

### Multiple sequence alignment and phylogenomic analysis of diatom myosin sequences

To identify diatom myosin sequences, the head domain of *C. australis* myosin A (CaMyoA) was used to conduct a BLAST search (E-value ≤10^−15^) against available diatom genome and transcriptome databases^61–63^. The retrieved protein sequences were manually curated to remove duplicates and include only those that contain motifs and domains that are indicative of a myosin head domain (i.e. P-loop, switch-I region, acting binding site and IQ motifs). A multiple sequence alignment of the head domains (sequence up to the first IQ motif) was performed in Clustal Omega with the default parameters. The output PHYLIP alignment file was used for a maximum likelihood phylogenetic analysis using IQ-TREE v1.6.12^64^. To assess branch supports, the analysis employed 1000 ultrafast bootstrap replicates^65,66^ and 1000 replicates of the SH-aLRT algorithm^67^. The resulting tree file was visualized using the R package ggtree^68^ and nodes satisfying the conditions of UFBoot ≥95 and SH-aLRT ≥80 were highlighted.

### Confirmation of myosin gene structure

The full-length gene models of *C. australis* CaMyoA, CaMyoB, CaMyoC and CaMyoD were confirmed using RACE and RT-PCR. *C. australis* cDNA and gDNA were prepared as previously described^27^. Briefly, two nested RACE PCRs (primer combinations listed in **Table S1**) using cDNA attached to oligo (dT)25 Dynabeads (ThermoFisher Scientific) as a template. The resulting PCR products were cloned into the pJet1.2 (ThermoFisher Scientific) or pGEMT-easy (Promega), transformed into DH5α *E. coli* and the plasmid DNA was sequenced. Intron/exon boundaries were confirmed by performing a PCR using both cDNA and gDNA as a template, Q5 DNA polymerase (NEB) with primer pairs covering the entire predicted coding region (primer combinations listed in **Table S1**).

### Plasmid construction of GFP-fusion proteins

The final DNA sequence of all plasmids was confirmed by DNA sequencing.

#### pCa_rpl44_GFP

The eGFP gene was amplified by PCR using the sense primer 5’-ATTC**TAC GTA**<u>GCATGC</u>TCTAGAATGGTGAGCAAGGGCGAGGAG-3’ (SnaBI site bold, SphI site underlined, XbaI site italics) and the antisense primer 5’-ATTC**GCGGCCGC** TTACTTGTACAGCTCGTCCATG-3‘ (NotI site bold). The resulting PCR product was digested with SnaBI and NotI and cloned into the SnaBI and NotI sites of pCa_rpl44^27^. The resulting plasmid was termed pCa_rpl44_GFP.

#### CaMyoA

The CaMyoA gene was PCR amplified from gDNA using the sense primer 5’-ATAC **TACGTA**ATGGAAGATATCAAGTCCACAG-3’ (SnaBI site bold) and the antisense primer 5’-GTTCCCTCTCC TCCTACTTTTCA**TCTAGA**ATTC-3’ (XbaI site bold). The resulting PCR product was digested with SnaBI and XbaI and cloned into the SnaBI and XbaI sites of pCa_rpl44_GFP. The resulting plasmid was termed pCa_rpl44_CaMyoAGFP.

#### CaMyoB

The CaMyoB gene and promoter sequence (1,000 bp) were PCR amplified from gDNA using the sense primer 5’-GGC GAA TTG GGT ACC GGG CCC CCC TCG AGC ATT CAG CTG AAT GAA GGA TTT CAT G-3’ and antisense primer 5’-CTT GCT CAC CAT AGC TAC GGA TTC GGC AGC-3’. The eGFP Gene was PCR amplified from pCa_fcp_GFP^27^ using the sense primer 5’-CGA ATC CGT AGC TAT GGT GAG CAA GGG CGA G-3’ and the antisense primer 5’-CGT CCC ATG AAC TTT ACT TGT ACA GCT CGT CCA TG-3’. The CaMyoB terminator sequence (500 bp) was PCR amplified from gDNA using the sense primer 5’-CTG TAC AAG TAA AGT TCA TGG GAC GCC ACC-3’ and the antisense primer 5’-CAC AAC AGA ACA GAC GGA CTG CAG GAG CAG GCT CAG CTG CGT G-3’. The pCa_rpl44/nat plasmid^27^ was digested with XhoI and EcoRI, and was assembled with the PCR fragments in NEBuilder® HiFi DNA assembly reaction (NEB) and transformed into DH5α *E. coli*. The resulting final plasmid was termed pCaMyoBGFP+ Ca_rpl44/nat.

#### CaMyoC

The CaMyoC gene and promoter sequence (1,000 bp) were PCR amplified from gDNA using the sense primer 5’-GGC GAA TTG GGT ACC GGG CCC CCC TCG AGG GCA CTC CTG GCA ACA TTC GTA CTA G-3’ and antisense primer 5’-CTT GCT CAC CAT CAG GCC TGG GGC GGA TGC −3’. The eGFP Gene was PCR amplified from pCa_fcp_GFP^27^ using the sense primer 5’-CGC CCC AGG CCT GAT GGT GAG CAA GGG CGA G-3’ and the antisense primer 5’-CTG CTG CCG TCG CTT ACT TGT ACA GCT CGT CCA TG-3’. The CaMyoC terminator sequence (500 bp) was PCR amplified from gDNA using the sense primer 5’-CTG TAC AAG TAA GCG ACG GCA GCA GTA TTT TC-3’ and the antisense primer 5’-CAC AAC AGA ACA GAC GGA CTG CAG GCA GAT CTG CAC CAA GAA GAA C-3’. The pCa_rpl44/nat plasmid^27^ was digested with XhoI and EcoRI, and was assembled with the PCR fragments in NEBuilder® HiFi DNA assembly reaction (NEB) and transformed into DH5α *E. coli*. The resulting final plasmid was termed pCaMyoCGFP+ Ca_rpl44/nat.

#### CaMyoD

The CaMyoD gene was PCR amplified from gDNA using the sense primer 5’-AAA CAA CAA TTA CAA CAA CCT ACA TGC CAA AGG AAA AGGAC −3’ and the antisense primer 5’-CCT CGC CCT TGC TCA CCA TTC TAG AAT CGG AAT CAG AGT CCG AG −3’. The pCa_fcp_GFP plasmid^27^ was digested with SnaBI and XbaI, and was assembled with the PCR fragments in NEBuilder® HiFi DNA assembly reaction (NEB) and transformed into DH5α *E. coli*. The resulting plasmid was termed pCa_fcp_CaMyoDGFP.

#### Actin

The Ca_rpl44 promoter sequence was PCR amplified from the plasmid pCa_rpl44^27^ using the sense primer 5’-ATT GGG TAC CGG GCC CCC CCG TCC GTC TGT TCT GTT GTG-3’ and the antisense primer 5’-TGC TCA CCA TGT ACT TTG CAA TTC AAT TCA AGA TAC-3’. The eGFP Gene was PCR amplified from pCa_fcp_GFP^27^ using the sense primer 5’-TGC AAA GTA CAT GGT GAG CAA GGG CGA G −3’ and the antisense primer 5’-CGT CAG ACA TAC CAC CAC CAC CCT TGT ACA GCT CGT CCA TGC-3’. The *C. australis* actin gene was PCR amplified from gDNA using the sense primer 5’-GGT GGT GGT GGT ATG TCT GAC GAC GAA GAT ATC-3’ and antisense primer 5’-TTG CGC GGC CTT AGA AGC ACT TGC GGT G-3’. The Ca_rpl44 terminator sequence was PCR amplified plasmid pCa_rpl44^27^ using the sense primer 5’-GTG CTT CTA AGG CCG CGC AAA CCA AAC G-3’ and the antisense primer 5’-CAG AAC AGA CGG ACT GCA GGA GCT CCG ATG GCA ACA GC-3’. The pCa_rpl44/nat plasmid^27^ was digested with XhoI and EcoRI, and was assembled with the PCR fragments in NEBuilder® HiFi DNA assembly reaction (NEB) and transformed into DH5α *E. coli*. The resulting final plasmid was termed pCa_rpl44_GFP_gly4__actin + Ca_rpl44/nat.

### Genetic transformation of *C. australis*

Biolistic transformations to introduce genes for GFP fusion proteins were performed using a previously published protocol^27^. Briefly, 1 × 10^8^ wild type cells were plated on a 5 cm circle in the middle of a 1.5 % EASW agar plate. Tungsten particles coated with plasmid DNA were then delivered into the cells using the Biorad PDS-1000/He™ Biolistic Particle Delivery System (28 mmHg vacuum at 1550 psi rupture). Cells were scraped from the plate and grown in liquid culture medium for 24 hours and then 5 × 10^6^ cells were plated onto 1.5 % EASW agar plates supplemented with 450 µg·mL^−1^ nourseothricin. Single colonies became visible following 10-14 days and were transferred into liquid EASW medium supplemented with 450 µg·mL^−1^ nourseothricin.

### Spinning Disk/Laser Scanning Confocal Fluorescence Microscopy

The immobilization of cells used for confocal microscopy was achieved through one of two methods: i) 10 µL of a diatom cell suspension was pipetted on a glass-bottomed cell culture dish (Ibidi, Gräfelfing, Germany, Cat. No. 80827), and overlaid with a thin slice of 1 % (w/v) agarose in EASW, ii) 300 µL of cell suspension were pipetted into a glass-bottomed 8-well cell culture dish (Ibidi, Cat. No. 81156) and left to settle for 45 min at 18°C under constant light. The medium was then removed gently using a micropipette, and replaced with 300 µL of 1 % (w/v) low melting (gelling temperature: 25±5 °C) temperature agarose (Fisher Bioreagents, Cat. No. BP165) prepared in EASW.

Laser scanning and spinning disk confocal microscopy were at the Light Microscopy Facility and Molecular Imaging and Manipulation Facility respectively, both Core Facilities of the CMCB Technology Platform at TU Dresden. Laser scanning confocal fluorescence microscopy (LSCM) was performed using an LSM 780/FLIM inverted microscope (Zeiss, Jena, Germany) equipped with 32-channel GaAsP spectral detectors and using a Plan-Apochromat 63x/1.46 Oil Corr M27 (Zeiss) and the Zen software (2011 version, Zeiss). Images were captured using a simultaneous three channel mode to acquire fluorescence of GFP (excitation: 488 nm, emission: 489 - 524 nm), chloroplast autofluorescence (excitation: 633 nm, emission: 667/20 nm) and bright field images. Spinning disk confocal microscopy (SDCM) was performed using a Nikon Eclipse Ti-E inverted microscope equipped with an Andor iXon Ultra 888, Monochrome EMCCD camera and a 100x/1.49 SR Apochromat Oil Objective (NIKON) and NIS Elements software (version 4.5). Two laser lines were used to detect the fluorescence of GFP (excitation: 488 nm, emission: 525/30 nm) and chloroplast autofluorescence (excitation: 647 nm, emission: 685/40 nm).

### Total Internal Reflection Fluorescence (TIRF) Microscopy

TIRF Microscopy was performed on a Nikon Eclipse Ti2 microscope equipped with a perfect focus system (PFS), a 100X, 1.49 NA oil, apochromatic TIRF objective with either a 1x or 1.5x Optovar tube lens (pixel sizes: 130 x 130 nm and 86×86 nm respectively, with 1:1 aspect ratio). Samples were illuminated with a 488 nm laser placed in a visitron laser box and channeled through an iLas2 ring TIRF module operated in ellipse mode (ring TIRF). Images from different emission channels were acquired with separate EMCCD cameras (iXon Life EMCCD for 525/30nm (GFP) and iXon Ultra EMCCD for ≥653 nm (chloroplast) channel) each containing 1024 x1024 pixel sensor and controlled with VisiView software. Images were acquired in time-lapse mode every 100 ms with 100 ms exposure (10 frames per second). Movies were acquired for a total time of between 1-2 minutes.

### Data processing and analysis

#### Movies from the VisiView software were converted into .tiff format and processed using Fiji & MATLAB. Fiji commands and plugins used are shown in *italics*

##### Tracking

Regions of movies covering paths of single cells were cropped out either by defining a rectangular region of interest (ROI) and using *crop* or *duplicate*, or by drawing a line ROI covering the entire path followed by the cell, and then using the *straighten* command. Far-red channel data (showing chloroplast autofluorescence) was used together with *TrackMate*^69,70^ to obtain the X-Y positions over time, for a chloroplast signal in the leading and trailing half of the cell (**Supplementary** Fig 2b). Spot sizes were set to match bounding circles with the diameter of the larger chloroplast (∼4.5-6.0 µm, cyan and magenta open circles, **Supplementary** Fig 2b). Custom MATLAB code was used to determine the angle of the long axis of the cell with respect to the lab coordinate system using the vector connecting the two bounding circles. The center position of the cell (red filled circle) over time was defined as the geometric mean between the center positions of the two bounding circles (cyan and magenta filled circles in **Supplementary** Fig 2c). Instantaneous velocity values were obtained by dividing the frame to frame displacement of the cell’s center point by the time interval between consecutive frames (100 ms). All velocity graphs were generated in MATLAB and smoothed using the “smooth” function with a moving average of 20 data points to reduce noise caused by the time resolution of frame to frame tracking.

##### GFP-channel registration

A custom macro was written in Fiji to register GFP channel data. As a reference, a frame was selected in which the cell was oriented horizontally and located approximately in the center of the field of view (FOV). With respect to the orientation of the cell in this frame, all remaining frames were registered by *translate* and *rotate*, using the positions and angles calculated from the chloroplast channel as described above.

##### Kymograph generation

Kymographs of each horizontal row of pixels in registered GFP-channel stacks were generated using the *reslice* command (default parameters). These kymographs were then combined by generating a maximum projection via the *Z Project…* command in the Stacks menu. Combining these kymographs allowed us to better visualize the paths of fluorescent entity across the width of the cell (Supplementary S1d). Interference patterns and sudden changes in the z-position of the cell sometimes generated horizontal stripes of varying intensity, unrelated to the movement of fluorescently labeled entities. We removed these stripes using Fourier filtering. We performed a fast Fourier transformation of the image (FFT), assigned intensity values of 0 to a 2 pixel wide mask covering the height of the transformation, and back transformed it using *Inverse FFT*.

##### Myosin velocity measurements

We employed two different methods to measure the velocities of myosin spots from kymographs. For kymographs containing a large number of spots (with similar slopes), the *Directionality* plugin was used. To do so, we selected kymograph regions in which the cell showed smooth, sustained gliding (>0.5 µm·s^−1^ for at least 5 s). Rectangular ROIs (of 5-15 s) in the leading and trailing halves were drawn separately, such that they cover the paths of myosin spots (**Supplementary** Fig 2f**, indigo, red, cyan and yellow rectangles**). For each ROI, the *Directionality* plugin (method: Fourier Components) was used to generate a histogram of orientations of the paths of myosin spots. We used one of three values to estimate the mean myosin velocities in the ROI: (I) the peak of the Gaussian distribution of angles determined by the directionality plugin (most common), (II) the local maximum in the histogram of angles corresponding the paths of myosin (if the goodness of fit is poor), or (III) the manually measured angles of traces identified in the orientation map (if only a small portion of the predicted angles fit the traces in the kymograph). For kymographs containing a low number of spots (with variable slopes) velocities for straight segments were estimated using the *Straight Line* tool. The orientation of each segment was used to calculate myosin spot velocity. The above calculated velocities were then plotted against the average cell velocity calculated for the corresponding time window (**Fig. 4a**).

### Transmission Electron Microscopy

A culture of *C. australis* cells in a logarithmic stage of growth was pelleted through centrifugation (2000 xg, 2 min, 18°C) and then resuspended in a minimal volume of EASW. The concentrated cell suspension was filled into a high-pressure freezer copper/gold carrier (depth of 100 µm) pre-coated with a 0.1% lecithin in chloroform solution (M. Wohlwend GmbH, Switzerland). The loaded carriers were rapidly frozen under high pressure (Leica EM ICE High Pressure Freezer) using liquid nitrogen, and freeze substitution (Leica AFS2 automatic freeze substitution) was performed by immersing the specimens in a 1% osmium tetroxide solution in acetone for a duration of 70 hours. The substitution medium was subsequently replaced with pure acetone, and this step was repeated four times at 0°C. Following the substitution process, the samples were post-stained with 1% uranyl acetate in acetone for 1 hour on ice. Subsequently, they underwent multiple washes in acetone. To prepare the specimens for embedding, they were infiltrated with a 3:1 mixture of acetone and epoxy resin for one hour. This was followed by an overnight incubation in a 1:1 mixture at refrigerator temperature. The specimens were gradually allowed to reach room temperature before being further infiltrated with a 1:3 mixture of acetone and resin for 4 hours and an additional 1:3 mixture overnight at room temperature. Finally, the samples were incubated in pure epoxy resin for 4 hours and overnight. After the last resin change, the specimens were flat-embedded between Teflon-coated slides, and the resin was cured at 60°C for a duration of 48 hours. Ultrathin sections (70 nm) were cut using a diamond knife on an ultramicrotome and collected on formvar-coated copper slot grids. The sections were subsequently post-stained with uranyl acetate and lead citrate, following standard protocols^71–73^. The samples were evaluated using a Morgagni transmission electron microscope (FEI, Thermofisher) operating at 80 kV, and micrographs were acquired using a Morada CCD camera (EMSIS). This work was performed at the Electron Microscopy facility of the Max Planck Institute of Molecular Cell Biology and Genetics.

## Data Availability

The genomic DNA and protein sequences have been deposited at GenBank: CaMyoA: PP083301; CaMyoB PP083302; CaMyoC: PP083303; CaMyoD: PP083304 All raw data and code used to generate the figures in this paper can be found in the following Zenodo record: https://doi.org/10.5281/zenodo.10722975.

## Code Availability

Modifiable versions of Fiji and MATLAB code used to analyze microscopy data can be found in the following repository: https://github.com/metingd/DiatomMotilityMyosin. Code used to generate the phylogenomic tree can be found in the Zenodo record shown above.

## Acknowledgments

We thank Jennifer Klemm, Corina Bräuer and Martina Lachnit for technical help. This work was supported by the Light Microscopy Facility and the Molecular Imaging and Manipulation Facility, Core Facilities of the CMCB Technology Platform at TU Dresden, and the Electron Microscopy Facility at the Max Planck Institute of Molecular Cell Biology and Genetics Dresden and by grants from the Deutsche Forschungsgemeinschaft (PO 2256/1-1) to N.P., (KR 1853/9-1) to N.K. R.H. was supported by the Deutsche Forschungsgemeinschaft (DFG, German Research Foundation) under Germanýs Excellence Strategy – EXC-2068 – 390729961-Cluster of Excellence Physics of Life of TU Dresden.

## Contributions

NP, NK and SD conceived the project. NP, NK, SD, MGD, VFG, LN and RH conceptualized and designed the experimental research and analyses. MGD, LN and NP generated the protein constructs and performed the experiments. MGD, VFG, LN, MLZ and JRS wrote code for analyses and analyzed the data. MGD, NP, NK and SD wrote the manuscript. All authors discussed the data, reviewed and edited the manuscript.

## Corresponding authors

Correspondence to Nicole Poulsen and Stefan Diez

