## Supplementary Information for "Gliding motility of the diatom *Craspedostauros australis* correlates with the intracellular movement of raphid-specific myosins"

### Supplementary Figures

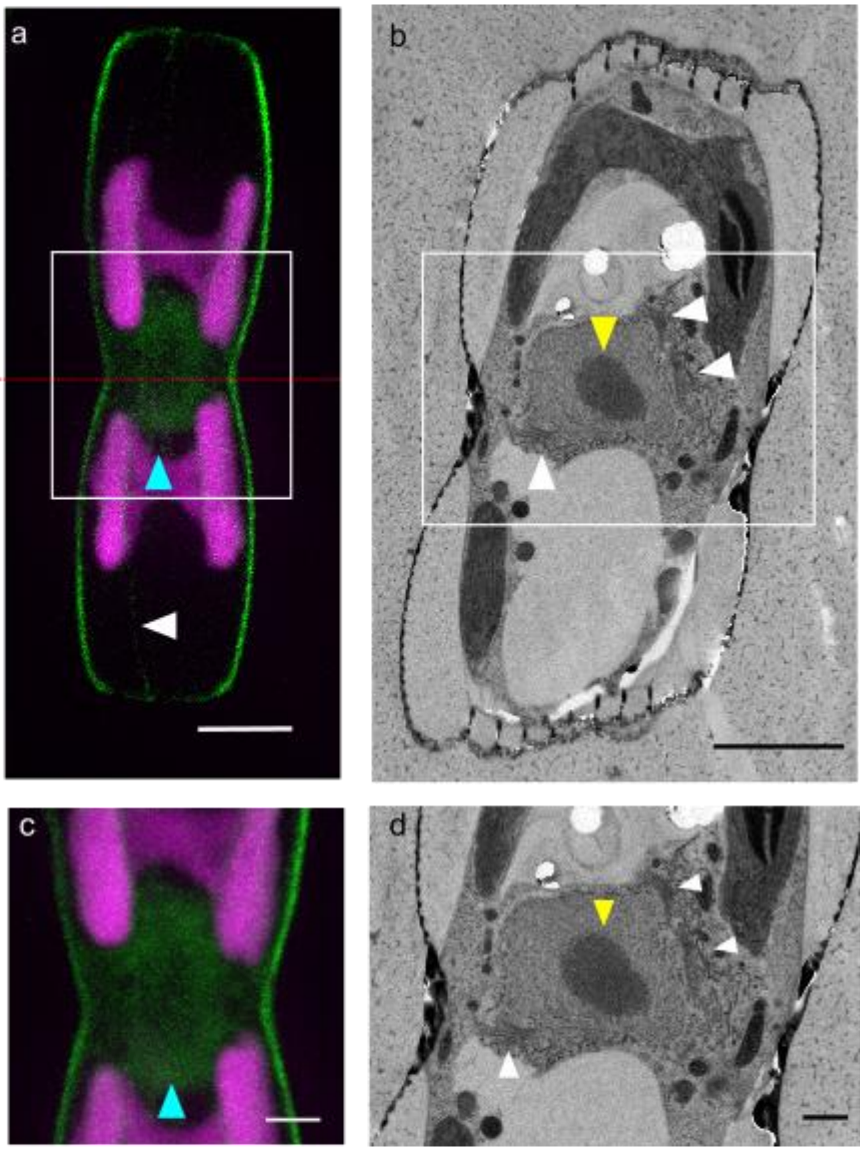


**Supplementary Figure 1: Confocal fluorescence microscopy of *C. australis* cells expressing GFP-actin and transmission electron microscopy (TEM) of sectioned wild type cells. a, c** Confocal fluorescence microscopy image showing a maximum projection of an entire cell (GFP-actin in green, chloroplast autofluorescence in magenta). **c** Region indicated by a white box in **a** showing the distribution of actin in the perinuclear region (cyan arrowheads) and actin along the girdle band (white arrow). **b, d** Transverse TEM cross section of a wild type cell close to the midpoint of the cell and the central nodule (approximate location indicated by red dotted line in **a**). **d** Region indicated by a white box in **b**. The Golgi complex (white arrowheads) can be seen in the area surrounding the nucleus (yellow arrowheads). Scale bars – a: 5 µm, b,c 2 µm, d: 0.5 µm.

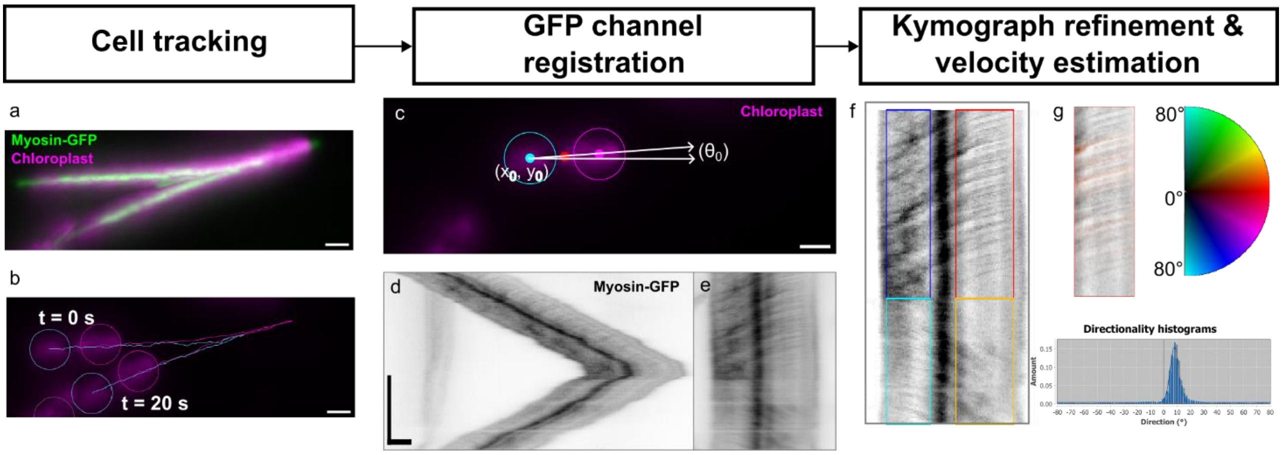


**Supplementary Figure 2: Image analysis pipeline. a** Maximum projection of a 20 s dual-channel movie of a gliding cell. **b** Overlay of tracks generated using TrackMate using the chloroplast autofluorescence signal in the far red optical spectrum. Chloroplast positions (circles) are shown only for the first and last frames. **c** Chloroplast positions for each frame were used to determine the position of the midpoint between the two chloroplasts (red dot) and the angle of the cell relative to a horizontal origin. **d, e** Kymographs of myosin-GFP signals (in black) generated by reslicing and projecting each row of pixels of the image before (**d)** and after (**e**) registration using displacement and angle values. **f** Refined kymograph after performing a Fast-Fourier-Transformation, masking horizontal traces, and inverting the Fourier transformation. Indigo, red, cyan and yellow boxes indicate regions-of-interest (ROIs) used for directionality measurements. **g** Orientation map showing the preferred orientation of the myosin-GFP signals in the red ROI shown in **f** (top left), color wheel showing gradient of colors used to indicate angle (top right) and histogram of estimated angles generated by the directionality plugin in Fiji. Scale bars – horizontal 5 µm, vertical 5 s.

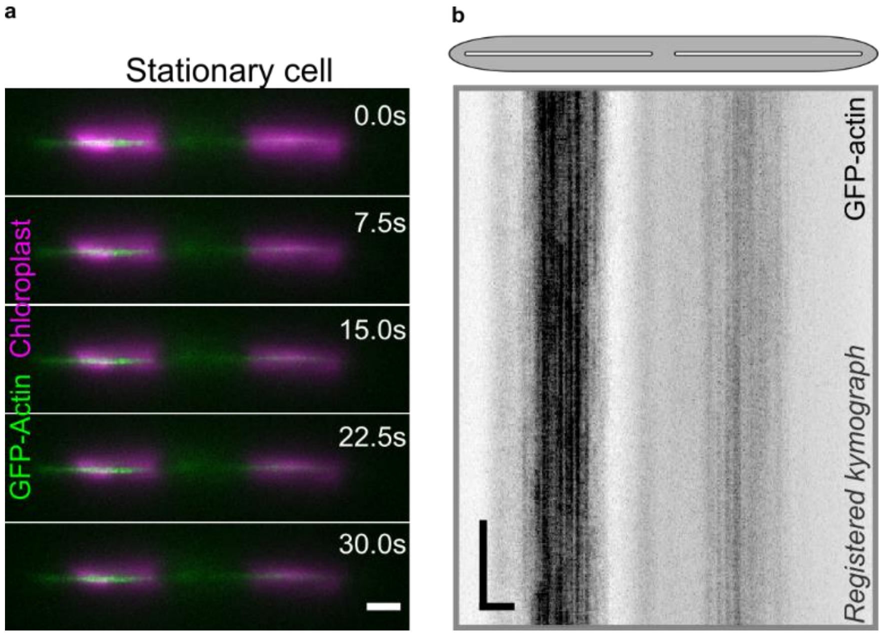


**Supplementary Figure 3: TIRFM based kymograph analysis of GFP-actin in stationary cells. a** Montage generated for dual-channel TIRFM data showing the positions of a stationary cell at 7.5 s intervals (GFP-actin in green and chloroplast autofluorescence in magenta, generated from Supplementary Movie S2). **b** Kymograph analysis of GFP-actin movement. GFP-channel data from the Supplementary Movie S2 shown in **a** was registered using cell tracking data (GFP-actin in black). Gray ellipse and white slits above kymograph approximate the positions of the cell body and raphe openings, respectively. (Scale bars: horizontal = 5 µm, vertical = 5 s)

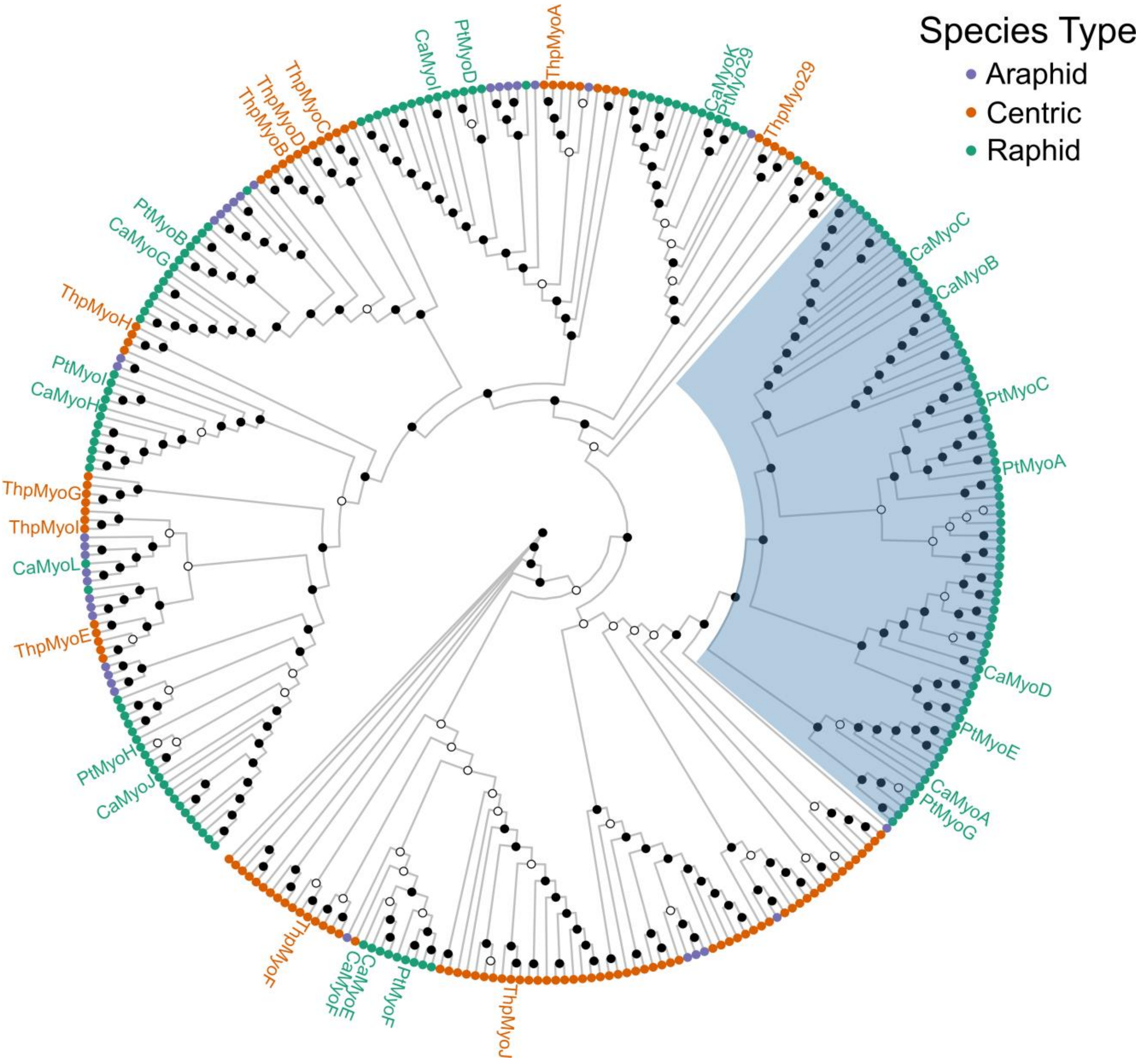


**Supplementary Figure 4: Phylogenomic analysis of diatom myosins.** Cladogram of 320 diatom myosins. The position of *C. australis* myosins (CaMyoX, 12 sequences), and previously published *Thalassiosira pseudonana* (ThpMyoX, 11 sequences) and *Phaeodactylum tricornutum* (PtMyoX, 10 sequences) myosins are labeled. A large clade containing only raphid diatom myosins, including the four *C. australis* myosins investigated in this study is highlighted in blue. Node tips are color coded by species type. Internal nodes indicate Ufboot bootstrap scores of <95 (hollow nodes) or ≥95 (solid nodes)

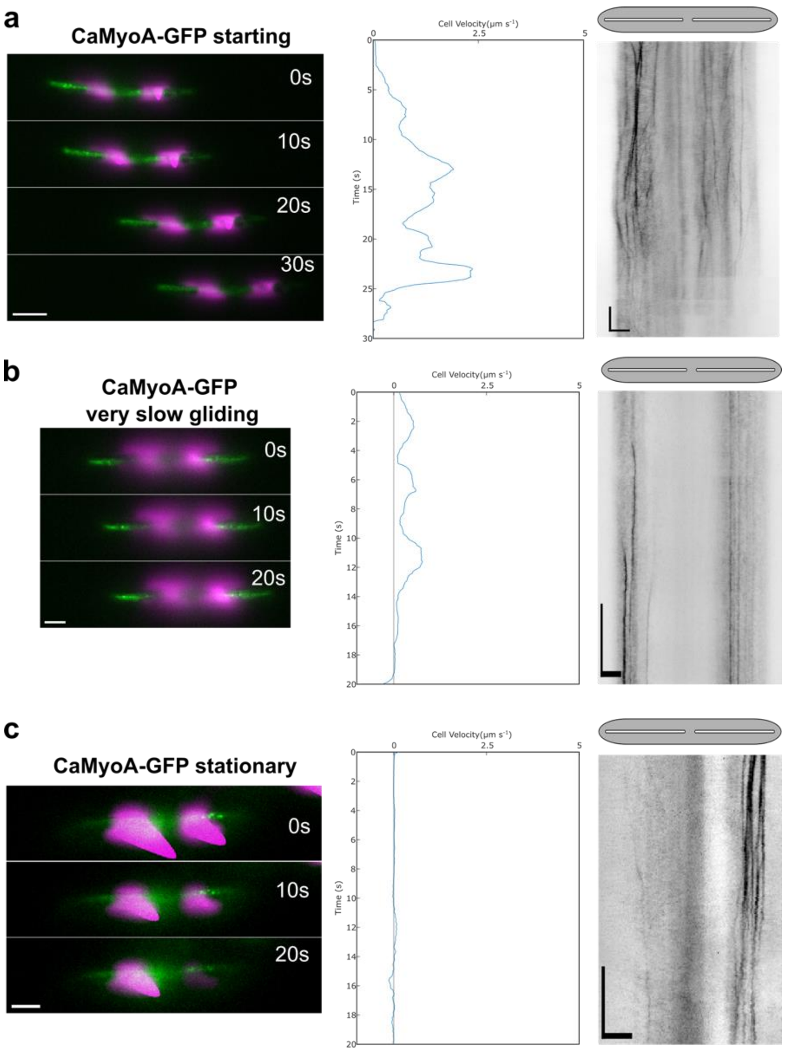


**Supplementary Figure 5: Kymograph analysis of CaMyoA-GFP in starting, slowly-moving and stationary cells.** Analysis of 25-30 s time-lapse segments of CaMyoA-GFP expressing cells upon **a** starting (Supplementary Movie S7), **b** slowly-moving (Supplementary Movie S8) and **c** being stationary (Supplementary Movie S9). (Left panels) Montages showing the position of cells at 10 s intervals (GFP in green, chloroplast autofluorescence in magenta, scale bars: 5 µm). (Middle panels) Cell velocity as function of time, generated from chloroplast tracking data. (Right panels) Registered kymographs generated from GFP-channel data (black) showing movement of myosins relative to the cell. Gray ellipses and white slits above kymographs approximate the positions of the cell body and raphe openings, respectively. (Scale bars: horizontal = 5 µm, vertical = 5 s)

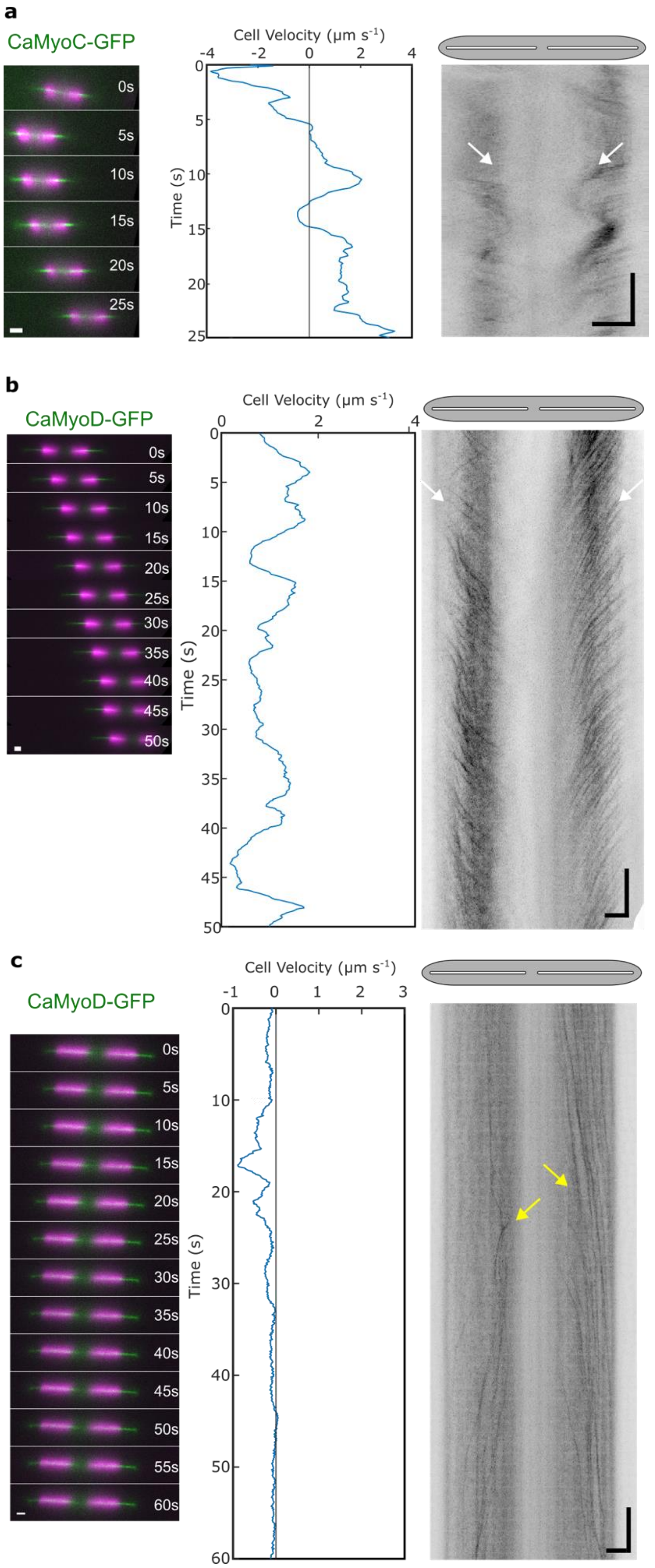


**Supplementary Figure 6: Multi-directional movements of CaMyoC-GFP and CaMyoD-GFP.** Analysis of 25-60 s time-lapse segments of CaMyoC-GFP and CaMyoD-GFP expressing cells showing **a** CaMyoC-GFP movement from apices to the cell center (white arrows) in a cell exhibiting repeated changes in gliding velocity, **b** CaMyoD-GFP movement from apices to the cell center (white arrows) in a cell exhibiting slow unidirectional gliding, and **c** CaMyoD-GFP movement from the center to the cell apices (yellow arrows) in a mostly stationary cell. (Left panels) Montages showing the position of cells at 5 s intervals (GFP in green, chloroplast autofluorescence in magenta, scale bars: 5 µm). (Middle panels) Cell velocity as function of time, generated from chloroplast tracking data. (Right panels) Registered kymographs generated from GFP-channel data (black) showing movement of myosins relative to the cell. Gray ellipses and white slits above kymographs approximate the positions of the cell body and raphe openings, respectively. (Scale bars: horizontal = 5 µm, vertical = 5 s)

### Supplementary Table

#### Table S1: Primers used to determine full-length gene models

| **Gene** | **Sequence** |
|---|---|
| **CaMyoA** | |
| 3'RACE | 1ST PCR: 5'-AGGGTCCTCGATTCAAACCCGCTCCTCGAAGCA-3';<br>2nd PCR: 5'-GTGCGTCCGAAGGGCTCCGACAA-3' |
| 5'RACE | 1ST PCR: 5'-CAGGGTACCCTATTCGAAACCGAAGATGTCGAG-3';<br>2nd PCR: 5'- GCCAGC GCGGATGCTTCGTAGA-3' |
| Full-length gene model | sense: 5'-CCACAGACAGCGCTGCGACT-3'<br>antisense 5'-CAGGGTACCCTATTCGAAACCGAAGATGTCGAG-3' |
| Full-length gene model | sense: 5'-AGGGAATTCGATTCAAACCCGCTCCTCGAAGCA-3'<br>antisense: 5'-GGC GATGTAGGTTGCCTT-3' |
| **CaMyoB** | |
| 3'RACE | 1st PCR: 5'-GCAGCCAGATCATCGCTTACC-3'<br>2nd PCR: 5'-CGCAAGGACTTCGACATCATG-3' |
| 5' RACE | 1st PCR: 5'-CCCTCGTGGTCCTCCCAGACG-3'<br>2nd PCR: 5'-GGGCAAGTTGACCATGTCCGG -3' |
| Full-length gene model | sense: 5'-AAACAACAATTACAACAACCTACATGGGCAAGAAAAAGGCC-3'<br>antisense: 5'- CCTCGCCCTTGCTCACCATTCTAGAAGCTACGGATTCGGCAGC-3' |
| **CaMyoC** | |
| 3'RACE | 1st PCR: 5'-CCTCATCCAGGCCAACGAGTC-3'<br>2nd PCR: 5'-CGTTCGAGATGTTGAACAACC-3' |
| 5' RACE | 1st PCR: 5'-GCTCCACCATGTCTGGGAAGG-3'<br>2nd PCR: 5'-ACTGTGGCACGCGGACTTGGG-3' |
| Full-length gene model | sense: 5'-AAACAACAATTACAACAACCTACGTATGGGCGAGAAAAAGTCCCAATACG-3'; antisense: 5'-CCTCGCCCTTGCTCACCATTCTAGACAGGCCTGGGGCGGATGC-3' |
| **CaMyoD** | |
| 3'RACE | 1ST PCR: 5'-TGCGAGAGCATGAAGCGTGAC-3'<br>2nd PCR: 5'-CGCGAAGGGATTGAACGCAAC-3' |
| 5' RACE | 1st PCR: 5'-GATACGGTTGACGACCGCATC-3'<br>2nd PCR: 5'- GTCGCCGACGTTCTTGGGCTC-3' |
| Full-length gene model | sense: 5'-AAACAACAATTACAACAACCTACATGCCAAAGGAAAAGGAC-3'<br>antisense: 5'- CCTCGCCCTTGCTCACCATTCTAGAATCGGAATCAGAGTCCGAG-3' |

### Supplementary Movies

| |
|---|
| Supplementary Movie 1 TIRFM of gliding GFP-actin expressing cell |
| Supplementary Movie 2 TIRFM of stationary GFP-actin expressing cell |
| Supplementary Movie 3 TIRFM of smooth, sustained gliding CaMyoA-GFP expressing cell |
| Supplementary Movie 4 TIRFM of smooth, sustained gliding CaMyoB-GFP expressing cell |
| Supplementary Movie 5 TIRFM of smooth, sustained gliding CaMyoC-GFP expressing cell |
| Supplementary Movie 6 TIRFM of smooth, sustained gliding CaMyoD-GFP expressing cell |
| Supplementary Movie 7 TIRFM of transition from stationary to gliding of CaMyoA-GFP expressing cell |
| Supplementary Movie 8 TIRFM of CaMyoA slow cell - more activity |
| Supplementary Movie 9 TIRFM of CaMyoA stationary cell - most activity |
| Supplementary Movie 10 TIRFM of transition from stationary to gliding of CaMyoB-GFP expressing cell |
| Supplementary Movie 11 TIRFM of transition from gliding to stationary of CaMyoB-GFP expressing cell |
| Supplementary Movie 12 TIRFM of reversal event of CaMyoB-GFP expressing cell |
| Supplementary Movie 13 TIRFM of CaMyoC-GFP movement from apices to center of cell |
| Supplementary Movie 14 TIRFM of CaMyoD-GFP movement from apices to center of cell |
| Supplementary Movie 15 TIRFM of CaMyoD-GFP movement from center to apices of cell |

All movies of *C. australis* cells expressing GFP-fusion proteins of either actin or myosin were acquired by TIRFM. Movies show composites of GFP and chloroplast channels (upper panel), and registered movies of only the GFP channel data (lower panel). Scale bars: 5 µm.
